# Ground-state pluripotent stem cells are characterized by Rac1-dependent Cadherin-enriched F-actin protrusions

**DOI:** 10.1101/2023.06.15.544897

**Authors:** Shiying Liu, Yue Meng, Pakorn Kanchanawong

## Abstract

Pluripotent Stem Cells (PSCs) exhibit extraordinary differentiation potentials that can be propagated *in vitro* and thus are commonly utilized in regenerative medicine. While different types of PSCs exist which correspond to different stages of pluripotency during embryogenesis, important aspects of their biology in terms of their cellular architecture and mechanobiology have been less well understood, thereby limiting their tissue engineering application potentials. Since the actin cytoskeleton is a primary determinant of cell mechanical properties, here we sought to investigate how the actin cytoskeleton may be differentially regulated in different states of pluripotency. Comparing ground-state naïve mESCs and the corresponding converted prime epiblast stem cells (EpiSCs), we observed the actin cytoskeleton is drastically reorganized during the naïve to prime pluripotency transition. Strikingly, we reported a distinctive actin organization that appears to be unique to the ground state mESCs, whereby the isotropic cortical networks are decorated with prominent actin-enriched structures which contain cadherin-based cell-cell junctional components, despite not locating at cell-cell junctions. We termed these structures “cadheropodia” and showed they arise from the cis-association of E-cadherin extracellular domain. Our measurements revealed that cadheropodia is under low mechanical tension and exhibits minimal calcium dependence, consistent with its putative dependence on cis-interaction. Further, we identified Rac1 as a negative regulator of cadheropodia whereby active Rac1 induces its fragmentation and dissociation of β-catenin. Taken together, we described a novel actin-based structures in the ground-state mESCs, which may have potential roles in ground-state pluripotency and serve as useful structural markers to distinguish heterogeneous population of pluripotent stem cells.

## Introduction

Pluripotent stem cells (PSCs) are important cell types both as model systems for embryonic development (Trounson and DeWitt, 2016) and from the regenerative medicine / tissue engineering perspectives (Tabar and Studer, 2014). However, important aspects of PSC biology, especially their cellular architecture and mechanobiology, have been relatively underexplored. Recently, works by our group as well as others are beginning to highlight major mechanobiological differences between the PSCs and more commonly studied adult cell types (Bergert et al., 2021; De Belly et al., 2021; Narva et al., 2017; Stubb et al., 2019; Xia et al., 2019a), and as reviewed recently (Liu and Kanchanawong, 2022). For example, in our earlier work, using super-resolution microscopy we characterized the unique isotropic network architecture of the actin cytoskeleton observed in mouse Embryonic Stem Cells (mESCs) and elucidated the underlying regulatory mechanisms (Xia et al., 2019a). In conjunction, we and others have also observed that the actin cytoskeletal architecture of mESCs appears to be drastically remodelled with prominent stress fibre formation upon losing pluripotency (Boraas et al., 2016; Xia et al., 2019a). Consistent with this, mESCs are known to exhibit characteristic mechanorefractory properties (Chowdhury et al., 2010; Xia et al., 2019b; Y.-C Poh, 2010), while pluripotency exit is associated with a marked increase in mechanosensitivity (Verstreken et al., 2019), as well as a membrane-tension dependent decoupling of the plasma membrane from the actin cortex (Bergert et al., 2021; De Belly et al., 2021). Since the morphology and mechanical properties of most vertebrate cells are strongly governed by the architecture of the actin cytoskeleton (Fletcher and Mullins, 2010), these observations collectively imply important involvement by the actin cytoskeleton during different states of pluripotency.

PSCs can be either originally derived from pre– and post-implantation embryos (Brons et al., 2007; Evans and Kaufman, 1981; Martin, 1981; Tesar et al., 2007; Thomson et al., 1998) or from adult-cell reprogramming (Park et al., 2008; Takahashi et al., 2007; Takahashi and Yamanaka, 2006). By definition, PSCs are capable of differentiating into any specialized lineage of the three germ layers or trophoblasts (Martello and Smith, 2014; Xu et al., 2002). Additionally, self-organization of PSCs under appropriate micro-environmental control could recapitulate essential features of embryonic development (Sozen et al., 2018; Zheng et al., 2019). Importantly, it is now well appreciated that pluripotency is not a single unified state but rather an evolving landscape that spans from pre-implantation to post-implantation stage of development (Nichols and Smith, 2009; Shahbazi et al., 2017). Among PSCs, mouse embryonic stem cells (mESCs) derived from the inner cell mass (ICM) of developing pre– implantation blastocysts show unbiased naïve pluripotency, with unrestricted potential to give rise to all somatic cells and germ lines and can be easily incorporated into epiblast and re-enter embryonic development to produce chimeric embryos (Hackett and Surani, 2014; Weinberger et al., 2016). However, mouse epiblast stem cells (mEpiSCs) from post-implantation embryos, although still pluripotent with capability to differentiation into three germ layers, exhibit limited contribution to chimeras, and are thus called “primed pluripotency”. Correspondingly, these two distinctive pluripotent status display numerous morphological, transcriptional, and epigenetic differences (Atlasi and Stunnenberg, 2017). Interestingly, human ESCs, in spite of originating from ICM of pre-implantation embryo, resemble murine primed pluripotency (Hackett and Surani, 2014). Due to ethical issues, no equivalent derivations have been achieved in post-implantation human embryos. Thus, studies based on mouse ESCs and EpiSCs have been useful in furnishing important information on pluripotency transition in human embryonic development. In particular, mESCs can be converted *in vitro* into mEpiSCs by addition of Activin A and FGF2 (Guo et al., 2009; Rugg-Gunn et al., 2012), and thus can be highly amenable for obtaining molecular insights into the architecture and functions of PSCs in different pluripotent states.

Leukemia inhibitory factor (LIF), an activator of JAK-STAT3 pathways, has been widely used to support mESCs self-renewal and maintain their pluripotency in culture (Smith et al., 1988; Ying et al., 2008). Mouse ESCs conventionally cultured in serum containing medium supplemented with LIF (SMLIF) are generally heterogeneous and prone to fluctuation in their naïve status, attributable to the effects of serum upstream of a broad range of often conflicting signalling pathways (Hackett and Surani, 2014). In response, a defined culture formulation with two inhibitors (termed “2i”) including PD0325901 (an inhibitor of MAPK/ERK kinase (MEK) and ERK signalling) and CHIR99021 (a glycogen synthase kinase 3 (GSK3) inhibitor) supplemented in serum-free medium has been developed, which promotes an optimized state of naïve pluripotency in mESCs (Nichols and Smith, 2009; Silva et al., 2008; Silva and Smith, 2008; Ying et al., 2008). In addition to facilitating more homogeneous naïve pluripotency, mESCs in 2iLIF medium are also considered to be more similar to the “ground state”, which describes the unrestricted naïve pluripotent state established *in vivo* in the epiblast cells of mature blastocyst (Hackett and Surani, 2014; Nichols and Smith, 2009). Notably, “ground state” differs subtly from “naïve pluripotency”. Naïve pluripotency refers to the unbiased capacity to generate chimera embryos after blastocyst injection, no matter how exactly these cells reflect the developmental ground state built in vivo. In this case, both mESCs in SMLIF or 2iLIF are functionally naïve pluripotent as determined by chimera generation. However, mESCs in 2iLIF but not SMLIF can be regarded as “ground state” as 2iLIF cultures show reduced or absence of lineage-associated genes expression, closer to the E4.5 epiblast cells of blastocyst, according to single-cell transcriptomic analysis (Hackett and Surani, 2014; Nichols and Smith, 2009; Silva et al., 2008; Silva and Smith, 2008). Since the naïve mESCs in SMLIF condition have been characterized in our previous study (Xia et al., 2019a), here we primarily focus on ground-state mESCs in 2iLIF condition and the corresponding converted EpiSCs.

In this study, we sought to conduct a systematic investigation of how the architecture of the actin cytoskeleton is reorganized during the naïve to prime pluripotency transition using an *in vitro* model. Using structured illumination super-resolution microscopy, we observed a distinctive organization of the actin cytoskeleton that appears to be unique to the ground state (2iLIF) mESCs, whereby the isotropic cortical networks are decorated with prominent actin-enriched structures that co-localize with cell-cell junction components such as E-cadherin, α-catenin, β-catenin, and p120, but not cell-matrix adhesion proteins. Furthermore, these structures are not located at cell-cell junctional regions. These structures are termed “cadheropodia” to contrast with filopodial or lamellipodial protrusions which are commonly associated with integrin-based cell adhesions. We dissected the molecular mechanism governing the organization of cadheropodia, demonstrating that they arise from the association of the extracellular domain of E-cadherin, with the cytoplasmic domain of E-cadherin being surprisingly dispensable. Our biophysical characterizations showed that cadheropodia are under low mechanical tension and exhibit minimal dependence on calcium, which implicated the cis-interaction between E-cadherin ectodomains in its formation in contrast to the trans-interaction important for adherens junctions. Further, we identified Rac1 as a negative regulator of the formation of cadheropodia whereby, active Rac1 disassembles cadheropodia into smaller puncta and release β-catenin from cadheropodia. Taken together, we described a novel actin-based structures in the mESCs, which may have potential roles in ground-state pluripotency and serve as useful structural markers to distinguish heterogeneous population of pluripotent stem cells.

## Results

### Actin cytoskeleton reorganizes during naïve to prime pluripotency transition

To obtain more homogeneous naïve and ground state mESCs, we first adapted mESCs from SMLIF to 2iLIF which resulted in characteristic bright, domed and compact colonies with refractive edges as observed under phase contrast microscopy (FigS1a). Subsequently, we converted ground-state mESC in 2iLIF into prime EpiSC using Activin A and FGF2 in basic N2B27 medium as described earlier (Guo et al., 2009) (Fig S1b). As observed by immunofluorescence microscopy (Fig 1a and Fig 1b), the naïve markers Klf4 and Nanog are significantly reduced in converted EpiSC (cEpiSC), while prime marker Otx2 is increased dramatically. Moderated reduction of Oct4 in cEpiSC compared to naïve mESC is also observed, consistent with its role as the core pluripotency marker and continued requirement to maintain pluripotency (Nichols and Smith, 2009). Further, we performed RNA-Seq analysis comparing naïve mESc and cEpiSC (Fig S1c), which also confirmed the reduction of naïve genes transcripts and the increase in primed pluripotency genes following the conversion (Guo et al., 2009; Sang et al., 2018; Tesar et al., 2007).

**Figure 1.**
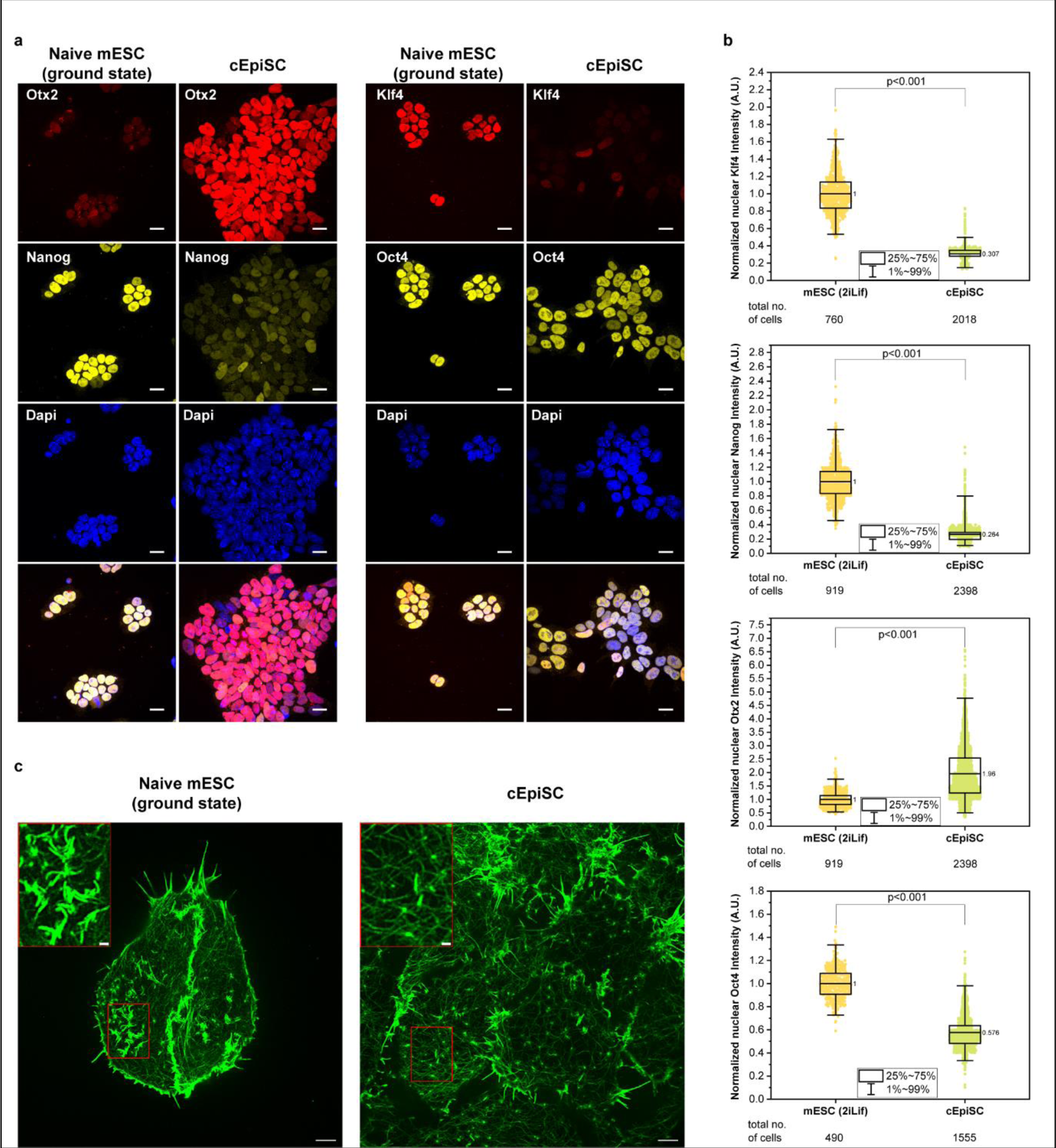
Actin cytoskeleton reorganizes during naïve to prime pluripotency transition and naïve ground state mESCs exhibit unique actin-rich structures. (**a**) Confocal micrographs of immunofluorescence staining for naïve mESC in 2iLIF ground state and converted epiblast stem cells (cEpiSC). Scale bar: 20 μm. **(b)** Box plots for quantitative analysis of the transcription factors in naïve mESC in 2iLIF ground state and cEpiSC. N=3 independent batches of experiments. The mean values were displayed. **(c)** F-actin structure imaged by spinning disk confocal-based structured illumination microscopy (SD-SIM) for naïve mESC and cEpiSC. Scale bar: 5 µm. Inset scale bar: 1 µm.

Earlier we have shown that distinctive actin cortical architecture can be probed in SMLIF mESC using spinning disk confocal-based structured illumination microscopy (SD-SIM) which provide ∼125 nm resolution (Xia et al., 2019a). To analyse the architecture of the actin cytoskeleton in 2iLIF ground state mESC and prime cEpiSC, we next applied SD-SIM to image their actin cytoskeleton as labelled by fluorescently-labelled phalloidin. As shown in Fig 1c and S1d, we observed that the actin cytoskeleton in both 2iLIF mESC and cEpiSC are organized into primarily low density cortical meshwork surrounded by lamellipodia and filopodia at cell edges and notable for the absence of stress fibers, similar to in SMLIF mESC reported earlier (Xia et al., 2019a) (note that as cEpiSC is known to suffer from low viability after replating as single cell (Kim et al., 2013), cEpiSCs were imaged as a colony). Interestingly, we also observed actin-rich structures in the interior of naïve 2iLIF mESC as shown in the inset images (Fig. 1c). These elongated structures appear to be excluded from the cell edge and strikingly disappear upon conversion into cEpiSCs, suggestive of their potential roles in naïve ground state pluripotency.

### Cadherin-dependent actin-rich structures are unique to ground-state mESCs

To investigate the molecular components of the actin-rich structures in ground-state mESCs, we next sought to inventory actin-binding proteins that may colocalize with these structures using immunofluorescence microscopy. Given the elongated appearance, one possibility is that these structures may represent a form of filopodia, a finger-like actin-rich protrusions that sense the surrounding extracellular matrix (ECM) or neighboring cells (Jacquemet et al., 2015). However, upon immunostaining for myosin X (Fig S2a), a classic marker of filopodia (Jacquemet et al., 2015; Jacquemet et al., 2019; Mattila and Lappalainen, 2008), we observed no colocalization (inset images) with the interior actin-rich protrusion but with the *bona fide* filopodia (white arrows) at the cell edge instead. Next we also probed for fimbrin, an actin bundling protein that serves as a marker for microvilli, actin-based protrusions of the apical plasma membrane found in epithelial cells (Ikenouchi et al., 2013; Revenu et al., 2004). However, as shown in Fig S2b, we observed no co-localization of fimbrin with the interior actin-rich protrusions in the ground-state mESCs, while the antibody specificity to fimbrin is validated by staining microvilli structures in MDCK epithelial cells (data not shown). In the previous study, we have identified cortical asters as key structures that maintain the actin cortex in SMLIF mESCs (Xia et al., 2019a). However, immunostaining for key components of asters such as Capping protein (CapZb) reveals the absence of co-localization (Fig S2c). Likewise, no co-localization was observed when cells were immunostained by a panel of antibodies for integrin-based adhesions (paxillin, vinculin, zyxin), actin associated proteins (cofilin, myosin IIA, tropomyosin3, spectrin), or clathrin coated pits (clathrin) (Fig S2c-i). Taken together, these results suggest that the actin-rich structures observed in the 2iLIF ground-state mESCs may be distinct from filopodia, microvilli, cortical aster, focal adhesions, and clathrin coated pits.

Surprisingly, we found these special structures in the ground-state mESCs exhibit strong colocalizations with E-cadherin, α-catenin, β-catenin, and p120 catenin (Fig 2a,d and 3a,d), which are common components of adherens junctions (AJ). E-cadherin mediates cell-cell interactions, through homophilic trans-ligation of the extracellular domain (EC) and through cytoplasmic connection of intracellular domain (IC) to β– and p120 catenin, and indirectly via β-catenin to α-catenin which can directly bind to the actin cytoskeleton (Bruser and Bogdan, 2017; Mege and Ishiyama, 2017). Strikingly, while the actin-rich structures in the 2iLIF ground-state mESCs contain the core components of AJs, they are not located at cell-cell junctions, and can also be observed in isolated single cells (insets in Fig 2a,d and 3a,d). To distinguish these structures from conventional filopodia that typically depend on integrin-based cell adhesions, we thus termed these structure “cadheropodia” to denote its cadherin-dependent organization. Notably, cadheropodia appears to be unique to the naïve ground state pluripotency since it cannot be detected upon conversion into cEpiSC (Fig 2c,d and Fig 3c,d), where enrichment of p120, β-catenin, α-catenin, and E-cadherin are largely confined to the cell-cell junctions.

**Figure 2.**
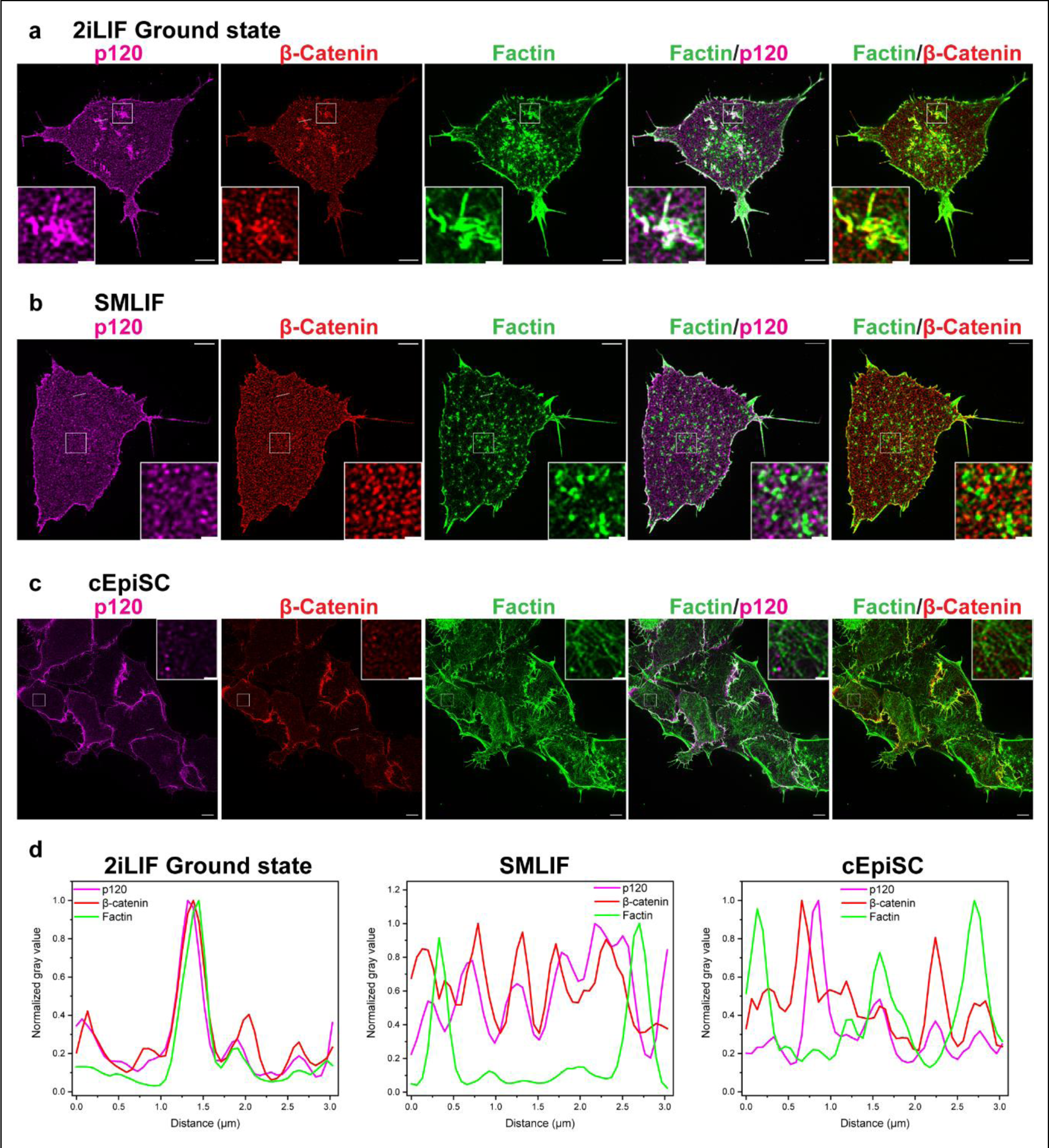
Actin-enriched cortical structures in 2iLIF ground state mESC cortex. (**a**) Confocal micrographs of immunofluorescence staining of p120 (magenta), β-catenin (red), and F-actin (green) for mESC in 2iLIF ground state **(a)**, mESC in SMLIF **(b)**, and cEpiSC **(c)**. Scale bar: 5 μm (inset: 1 μm). **(d)** Line scans of the corresponding channels through the line indicated in image **(a), (b)** and **(c).**

**Figure 3.**
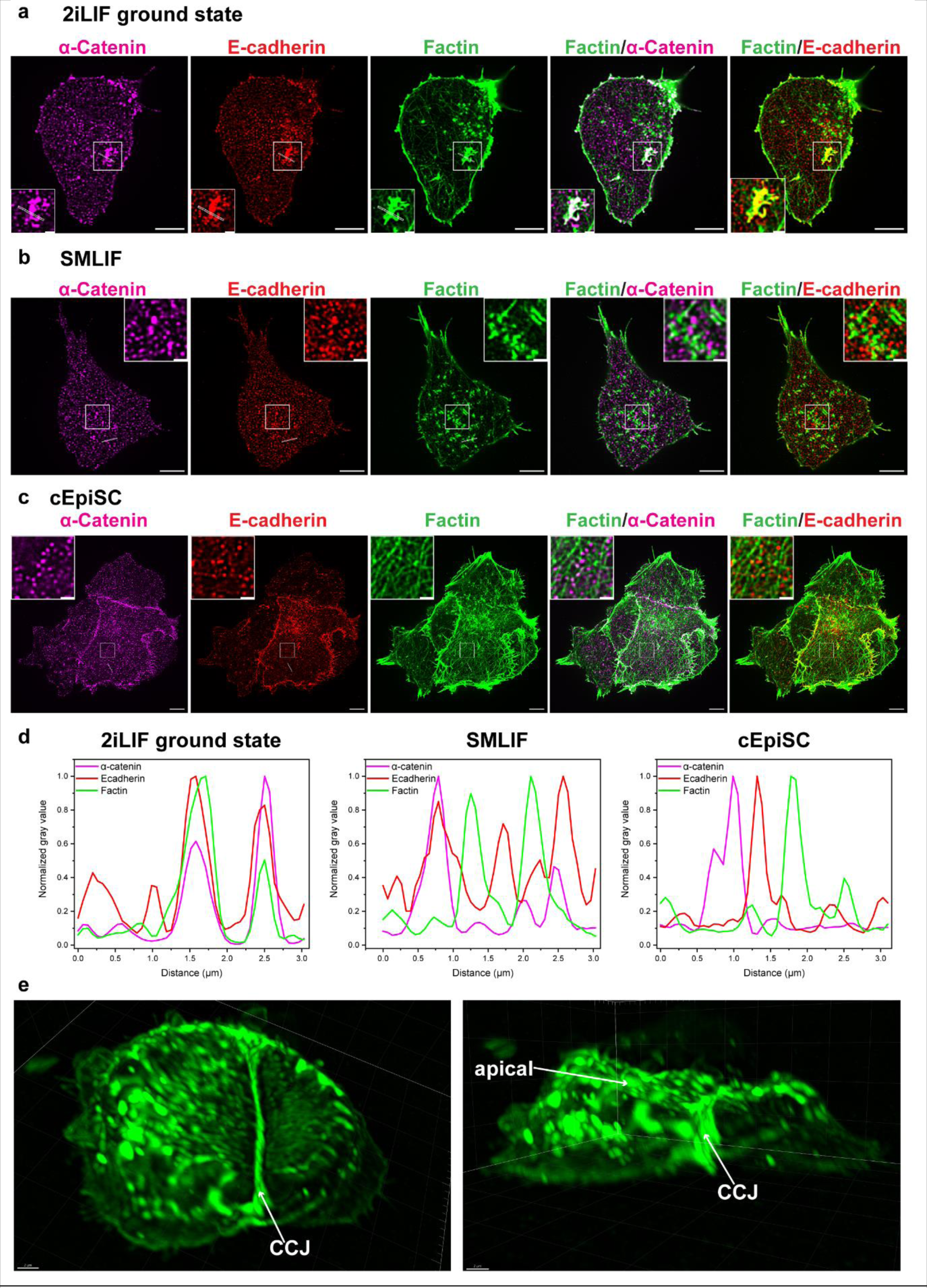
Cadheropodia colocalizes with α-Catenin and E-cadherin. (**a**) Confocal micrographs of immunofluorescence staining of α-Catenin (magenta), E-cadherin (red) and F-actin (green) for mESC in 2iLIF ground state **(a)**, mESC in SMLIF **(b)**, and cEpiSC **(c)**. Scale bar: 5 µm (inset: 1µm). **(d)** Line scans of the corresponding channels through the line indicated in image **(a), (b)** and **(c). (e)** 3D view images of E-cadherin GFP transfected mESCs in 2iLIF imaged by ZEISS Lattice Lightsheet 7 and exported by Imaris. CCJ: cell-cell junction.

Previous analysis by our group revealed that the actin cortex in SMLIF mESCs primarily consist of isotropic actin meshwork interspersed with Arp2/3 and α-actinin containing radial asters with putative roles in regulating cortical architecture (Xia et al., 2019a). As shown in (Fig 2b,d and 3b,d), immunostaining showed that the asters in SMLIF do not contain cadherin-catenin components (p120, β-catenin, α-catenin, and E-cadherin), nor does the cadheropodia of 2iLIF mESCs contain Arp2/3 complex and cadheropodia shows only partially colocalization with α-actinin (Fig S3a,b,c). Cadheropodia in 2iLIF ground state mESCs and asters in the SMLIF mESCs also differ in morphology and size, with the asters being typically circular and smaller than 0.4 um^2^ (Xia et al., 2019a). Quantitative analysis of cadheropodia using β-catenin immunostaining for segmentation showed that these structures are ∼0.5-0.6 um^2^ in area, and highly elongated and irregular with an aspect ratio ∼2.0-2.5, circularity of ∼0.20-0.23, and solidity of ∼0.50-0.55 (Fig S3d). To probe the dynamics of cadheropodia, we generated ground-state mESC with stable expression of LifeAct-mEmerald to enable live cell imaging. Here we observed that the cadheropodia is dynamic, with ∼30 min lifetime from appearance to disappearance (Fig S4a). Treatment with cytochalasin D disassembled cadheropodia into smaller puncta (Fig S4b), indicative of its dependence on actin polymerization. Interestingly, different cadheropodia structures respond with different dynamics to cytochalasin D treatment, (Fig. S4b: ROI 1 at ∼4min:36s, ROI 2 at ∼6min:35s and ROI 3 at ∼16min:14s after cytochalasin D addition). Cadheropodia also appears to be insensitive to myosin II contractility, as observed in the treatment with pNitro-Blebbistatin (supplementary movie 1), consistent with its lack of colocalization with myosin II (Fig S2g). Also, while most of our high-resolution imaging was performed for the ventral surfaces of the cells, upon using lattice lightsheet microscopy to observe mESCs transfected with E-cadherin-GFP, we also observed cadheropodia structure withinin the apical surface (Fig 3e). Altogether the observed differences in molecular components and morphodynamics suggest that cadheropodia and asters may be distinct cortical structures, and cadheropodia is a characteristic of the actin cortex in the 2iLIF ground-state mESCs.

### E-cadherin localization in cadheropodia depends on the cadherin extracellular domain

We next sought to understand the molecular basis of E-cadherin localization in the cadheropodia. E-cadherin mediates cell-cell interactions via the extracellular (EC) domain which contains five EC domain repeats (EC1-EC5) (Boggon et al., 2002). EC domains of E-cadherin participate in both trans– and cis-interaction, which respectively involve a strand swapping interactions of EC1 domains from opposing cells (Boggon et al., 2002), and EC1-EC2 interface of adjacent cadherin molecules (Harrison et al., 2011). Although cis-interaction is not required for AJ formation, it promotes the stability of cell-cell contacts through its effect on actin cytoskeletal anchoring to E-cadherin (Strale et al., 2015). The linkage between E-cadherin to the actin cytoskeleton is indirect, through adaptor proteins of the catenin family which bind to the intracellular domain (IC). Actin cytoskeletal connections are essential to the stability and lifetime of E-cadherin-mediated adhesions and also contribute to nanocluster organization of E-cadherin (Hong et al., 2013; Mege and Ishiyama, 2017; Wu et al., 2015).

To investigate the involvement of the EC and IC domains of E-cadherin, we expressed E-cadherin wild type (wt), mutants lacking EC (ΔEC) or IC domains (ΔIC) (Fig S5a) tagged with mScarlet-I into theground state 2iLIF mESCs. As observed by confocal microscopy in Fig 4a,b, wt E-cadherin primarily localized to cadheropodia (magenta arrows), basal cell-cell junctions (green arrows), apical cell-cell junctions (blue arrows), and intracellular vesicles as reported previously (Borghi et al., 2012). In contrast, E-cadherin lacking EC domain (ΔEC) failed to localize at the cadheropodia, as well as the basal and apical cell-cell junctions (Fig 4a,b). E-cadherin lacking intracellular domain (E-cadherin-ΔIC) appeard to localize substantially to both the cadheropodia (magenta arrow) and cell-cell junctions (green and blue arrows) as shown in (Fig S5b). These observations thus highlight the importance of EC domain in cadheropodia organization, consistent with its role in E-cadherin localization in other epithelial cells such as MDCK reported previously (Ozaki et al., 2010).

**Figure 4.**
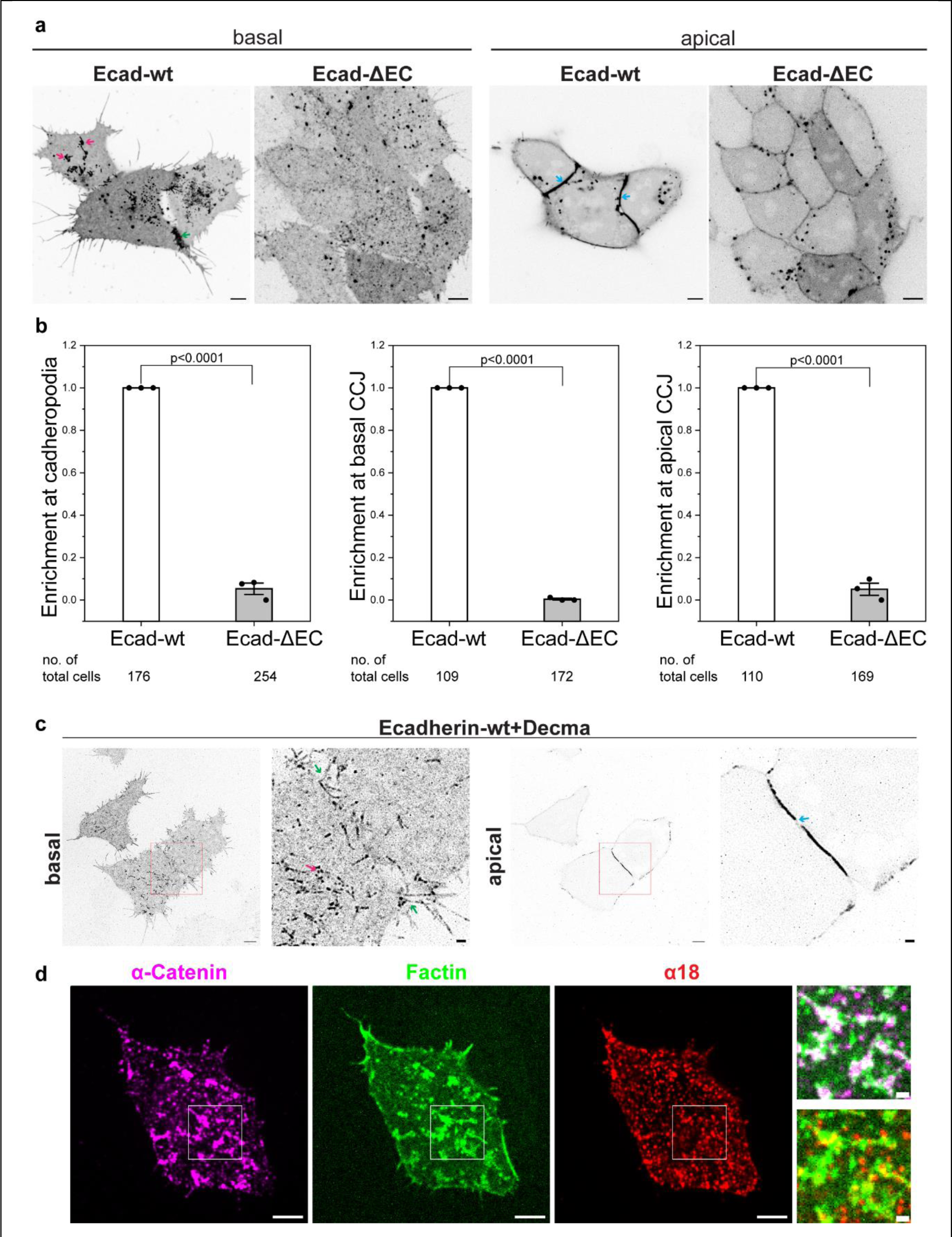
Extracellular domain of E-cadherin is essential for the localization at cadheropodia and cell-cell junction. (**a**) Confocal micrographs for live mESC in 2iLIF after transfection with E-cadherin-wild type (Ecad-wt) or E-cadherin without extracellular domain (Ecad-ΔEC) fused with mScarlet-I for 24h. Scale bar: 5 µm. (Red arrowhead: cadheropodia; green: basal CCJ; blue: apical CCJ). **(b)** Quantification for Ecadh-wt or Ecad-ΔEC enrichment at cadheropodia, basal CCJ, and apical CCJ from three batches of independent experiments. The percentages of cells with Ecad-ΔEC localization at cadheropodia, basal CCJ, and apical CCJ are normalized to those with Ecad-wt for every batch. Data is shown mean ± s.e.m. with total no. of cells shown at the bottom. Unpaired two-sample t-test is performed for statistics analysis with shown p value. **(c)** Live cell imaging for E-cadherin-GFP transfected mESC in 2iLIF after treatment with Decma antibody (10 µg/ml) for 17h.The magnified images in the red box are at right. Scale bar: 5 µm. inset scale bar: 1 µm. **(d)** Confocal micrographs for basal plane of mESC in 2iLIF after immunofluorescence staining with activated α-catenin (α18), α-catenin, and F-actin. Scale bar: 5 µm (inset: 1µm).

Next we perturbed inter-cadherin interaction using DECMA cadherin-blocking antibody which recognizes epitope in the EC domain (Brouxhon et al., 2013), observing that E-cadherin-wt localization became disjointed in the apical junction (blue arrow) (Fig 4c) and drastically diminished in the basal cell-cell junctions (green arrow indicating the cell-cell interface). We also observed a significant disruption of cadheropodia structures in the ventral plane, which exhibit isolated puncta features (magenta arrow) (Fig 4c). Based on a previous study showing that E-cadherin blocking antibody promotes E-cadherin endocytosis (Izumi et al., 2004), these structures likely correspond to incomplete cadheropodia structure that remained on the plasma membrane following DECMA antibody treatment.

### Cadheropodia is under low mechanical tension

Since cadherin-catenin complexes play important roles in mediating mechanical tension in AJs (Lecuit and Yap, 2015), we next sought to characterize whether the cadherin-catenin complexes in cadheropodia participate in mechanobiological roles. Laser nanoscissor has been used to ablate cellular structures to probe for mechanical tension from the analysis of initial recoil (Liang et al., 2016). As shown in (Fig. S5c and supplementary movie 2), when laser nanoscissor was applied to the apical junctions at the interface between mESCs in 2iLIF transfected with E-cadherin-GFP, significant recoil was observed, indicative of high level of junctional tension. On the contrary, the application of laser nanoscissor to the cadheropodia resulted in no obvious changes (Fig. S5d and supplementary movie 3), implying that cadheropodia may be under negligible level of tension. Consistent with this, we next probed the conformation of α-catenin in cadheropodia using α18 monoclonal antibody, which recognizes the mechanically-induced conformation of α-catenin that exposes vinculin binding site (Akira Nagafuchi, 1994; Yonemura et al., 2010). At the cell-cell junctions, clear α18 staining was observed as expected (Fig. S5e). In contast, as shown in Fig 4d, at the cadheropodia we observed the lack of α18 staining with non-conformation specific antibody staining (ab51032) suggesting that α-catenin at cadheropodia may remain in the mechanically inactive conformation, consistent with the lack of vinculin localization at cadheropodia (Fig. S2d).

### Cadheropodia is highly resistant to calcium depletion

Calcium binding at the interdomain linker regions of cadherin has been shown to promote E-cadherin rigidification and dimerization (Harrison et al., 2011; Nagar et al., 1996), while calcium chelator such as EGTA is commonly used to disrupt cell-cell junctions (Kartenbeck et al., 1991). To examine the calcium dependency of cadheropodia, 2iLIF mESCs expressing E-cadherin-GFP were treated with different concentrations of EGTA for specific time periods. Strikingly, as shown in Fig S6a and supplementary movie 4, a brief treatment (1 min) by 5 mM EGTA can dissociate apical junctions in mESC colonies (blue arrows), which continue to disaggregate over the next ∼30min. Interestingly, despite long 5 mM EGTA treatment of 1h (Fig S6b and supplementary movie 5) or even overnight (∼19h) (Fig S6c), cadheropodia appears to remain intact. Furthermore, cadheropodia remains abundant even after treatment with a high concentration of EGTA (20 mM) for 1h (Fig S6d and supplementary movie 6). These result suggests that cadheropodia is significantly more resistant to calcium depletion than AJs. Previously, it has been shown that the cis– and trans-interactions of E-cadherin depend on different concentrations of Ca^2+^ (Pertz et al., 1999), with low calcium (50 μM) able to stabilize crescent-like structure of E-cadherin molecule, medium concentration (500 μM) able to support cis-dimer formation, and high concentration (>1 mM) required for trans-interaction (Pertz et al., 1999). Our observation is thus consistent with the cis-interactions being the dominant organizing factor in cadheropodia, in contrast to trans-interaction which predominates in apical junctions.

### Constitutively active Rac1 disrupts cadheropodia organization

We next probe the roles of Rho family-GTPases, which are key regulators of cytoskeletal and cell adhesion dynamics, in the regulation of cadheropodia (Braga et al., 1997; Hall, 1998; Hodge and Ridley, 2016; Ridley and Hall, 1992; Ridley et al., 1992). We first expressed a panel of constitutively active (CA) forms of Rac1 (V12), cdc42 (Q61L), RhoA (V12) or dominant negative (DN) forms of Rac1 (N17), cdc42 (N17), Rho (N19) (Ridley et al., 1992; Wong et al., 2001) in 2iLIF ground state mESCs and observed their effects on cadheropodia. As shown in Fig 5a, the expression of CA-Rac1 drastically reduced the presence of cadheropodia (Fig 5b), with remaining actin-rich structures showing minimal co-localization with β-catenin, in contrast to control mESCs. On the other hand, the effect of CA-cdc42 expression depends on the expression level (Fig S7a), whereby cadheropodia was present under low level expression of CA-cdc42, but is largely suppressed and replaced by elongated filopodia at cell periphery upon high level expression of CA-cdc42. Meanwhile, the expression of CA-RhoA appears to minimally affect cadheropodia (Fig S7b), consistent with the absence of myosin IIA and low level of tension in cadheropodia (Fig S2g and S5d). The effects of DN Rac1, DN cdc42, and DN RhoA on cadheropodia appear to be minimal as transfected cells still contain prominent β-catenin-containing cadheropodia (Fig S7c,d,e). Taken together, these results suggest that the proper balance of Rac1 and, to a lesser extent, cdc42 likely participate in the regulation of cadheropodia.

**Figure 5.**
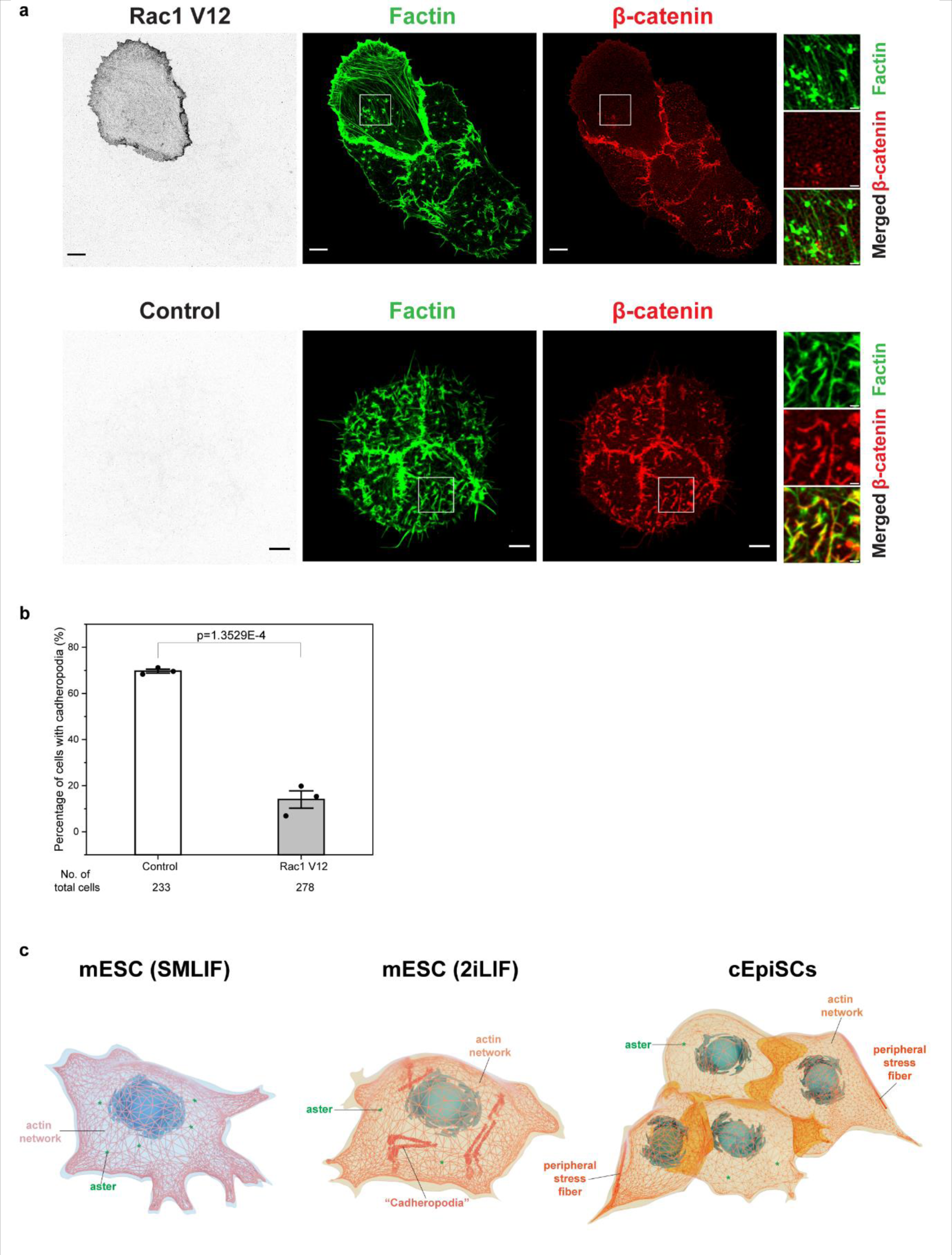
Active Rac1 disassembles cadheropodia. (**a**) Immunofluorescence images for control and Rac1 V12 transfected mESC in 2iLIF. Scale bar: 5 μm. Inset: 1μm**. (b)** Quantification for percentage of cells with cadheropodia in control and Rac1 V12 transfected mESC in 2iLIF from three independent experiments. Data is shown mean ± s.e.m. with total no. of cells shown at the bottom. Unpaired two-sample t-test is performed for statistics analysis with shown p value. (c) Schematic figure for comparing actin cortex of mESC in SMLIF, 2iLIF, and cEpiSC colony, highlighting the unique cadheropodia structure in mESC 2iLIF.

## Discussion

The actin cytoskeleton is capable of self-organization into a broad range of architecture to support different functions under different cellular contexts. While it has been observed that actin cytoskeletal remodelling occurs during embryonic stem cell differentiation, detailed understanding of these process has been less clear (Liu and Kanchanawong, 2022). Together with our earlier observation in naïve SMLIF mESCs (Xia et al., 2019a), in this study we observed that a common feature of the actin cytoskeleton in different pluripotent states of PSCs appears to be the baseline isotropic network in the cell cortex. As shown in the previous study, actin cortex of mESCs in SMLIF is at a low density with a large network pore size in the range of ∼350 nm, which effectively precludes contraction by myosin II minifilaments (∼300 nm). Thus, these cortical networks likely account for the low (<1kPa) elastic modulus of mESCs as well as their attenuated mechanoreponses (Chowdhury et al., 2010; Verstreken et al., 2019; Xia et al., 2019b; Y.-C Poh, 2010). On the other hand, localized actin-enriched structures on top of the cortex appear to be dependent on different pluripotent states. As summarized in Fig.5c, these include cadherin-enriched cadheropodia in ground-state pluripotent mESCs described in this study, the cortical asters analyzed earlier in naïve SMLIF mESCs (Xia et al., 2019a), and the peripheral stress fibers in EpiSCs (this study and (Amar et al., 2023; Narva et al., 2017; Rosowski et al., 2015)). Interestingly, these actin-based structures have been shown to perform functions related to the unique requirement of distinct pluripotent states. For example, the cortical asters in SMLIF mESCs are associated with the maintenance of sparse isotropic cortex that maintain the soft mechanics of mESCs, while the peripheral stress fibers in EpiSCs are thought to impart mechanical compression that maintain the pluripotency of the colony interior (Amar et al., 2023; Narva et al., 2017; Rosowski et al., 2015). As described above, the cadheropodia is observed primarily in the 2iLIF ground state but not in SMLIF mESCs. Furthermore, cadheropodia also disappears following conversion into primed pluripotent EpiSCs. Thus, one may surmise that potential roles of the cadheropodia could be as pre-assembled cadherin-catenin complexes that are poised to engage in cell-cell interactions upon the E-cadherin transligation, perhaps to enable rapid formation of cell-cell junctions. Indeed, it has earlier been shown that at the activated integrin primed for adhesion is positioned at the tip of filopodia, akin to ‘sticky fingers’ (Galbraith et al., 2007). At present, it remains challenging to dissect the function of cadheropodia, due to the technical challenge in selectively perturbing E-cadherin in the cadheropodia without affecting cell-cell junctions. In future investigation, the deployment of optogenetic approaches to spatiotemporally control cadherin activity (Mombo et al., 2023; Yu et al., 2020) or sub-cellular micromanipulation (Shakoor et al., 2022) may enable detailed functions of cadheropodia to be further elucidated.

Intriguingly, we also note that the cadheropodia observed here bears interesting parallel to E-cadherin-dependent filopodia observed in the 8-cell mouse embryo stage (Fierro-Gonzalez et al., 2013). Such E-cadherin filopodia protrude from the adherens junctions bordering the apical membrane of neighbouring cells and are necessary to maintain an elongated cell shape for the 8-cell embryo compaction (Fierro-Gonzalez et al., 2013). One notable difference, however, is the presence of myosin X in the cadherin filopodia at 8-cell stage (Fierro-Gonzalez et al., 2013), whereas myosin X is absent from the cadheropodia described in this study. Such differences potentially may reflect different stages of embryonic timeline, as in our study, the naïve ground-state to prime pluripotency transition can be considered to correspond to E3.5-4.5 pre-implantation blastocyst to E5.5 post-implantation embryo which is later than 8-cell embryo stage in the mouse (Atlasi and Stunnenberg, 2017). In recent years several unique cytoskeletal organizations have been described in early embryonic developments as well as in PSCs (Lim and Plachta, 2021; Liu and Kanchanawong, 2022). In this light, the cadheropodia may represent another example of unique sub-cellular architecture that arises during the specific stage of developmental morphogenesis. Indeed, the presence of cadheropodia and other pluripotent-state specific structural markers, may potentially serve as useful image-based markers for characterizing pluripotency state in heterogeneous population of cells.

Our observation of the cadheropodia also raises interesting questions of the molecular organization of cadherin in cells. Previous studies have shown that E-cadherin can be organized as clusters across different length scales on the cell surface regardless of cell-cell interactions (Borghi et al., 2012; Padmanabhan et al., 2017; Wu et al., 2015). For example, in MDCK cell model, in addition to cell-cell junction localization, exogenously expressed E-cadherin is found to exhibit occasional punctate-like clustering (Borghi et al., 2012). Non-junctional clusters of E-cadherin homologs were also observed in the *C. elegans* embryo zygote, which resist cortical flow and slow down cytokinetic furrow ingression by providing mechanical resistance to cortical deformations and decreasing RHO-1 and NMY-2 levels at the cortex (Padmanabhan et al., 2017). At the nanoscale, super-resolution microscopy has shown that E-cadherin is clustered into sub-100 nm domains both within and outside of adherens junctions, with domain size primarily determined by the cortical actin meshworks. The ectodomain of E-cadherin is known to be capable of both trans– and cis-interaction, which correspond to the interaction with the ectodomain of another E-cadherin molecule on an opposing cell or on the same cell, respectively (Brasch et al., 2012). From X-ray crystallography and computer simulation, the cis and trans interfaces are thought to be distinct, and thus a single cadherin can simultaneously engage in one trans– and two cis interactions, thereby promoting cluster formation (Harrison et al., 2011; Wu et al., 2011). The formation of AJ requires trans-interaction (Brasch et al., 2012), which can be readily disrupted by calcium depletion in contrast to cis-interaction, which is less susceptible to calcium (Pertz et al., 1999). In cells, cis interactions have been shown to be dispensable for AJ formation, as E-cadherin with mutations disrupting cis-interaction is still capable of forming AJ. However, the disruption impairs the stability of cell-cell contacts and coordinated behaviour in the collective cell migration (Strale et al., 2015). Instead, cis interactions contribute to the formation of ordered oligomers of E-cadherin, which help regulate anchoring of cadherins to the actin cytoskeleton and stabilize the junctions (Strale et al., 2015). In this context, our observation that calcium depletion, appears to minimally affect cadheropodia, suggests the E-cadherin clustering in cadheropodia may be relatively less susceptible to calcium. One possibility is that E-cadherin within the cadheropodia could be clustered through cis-interaction. Whether cadheropodia involves a close-packing organization of E-cadherin is currently unknown but could be addressed in future studies using E-cadherin with mutated cis interactions.

Thus far, we have identified Rac1 as being in control of the molecular pathways governing E-cadherin clusterings that give rise to cadheropodia. Overexpression of constitutively active Rac1 results in the disaggregation of cadheropodia into smaller puncta (Fig.5). Constitutive E-cadherin dimerization and subsequent clustering have been shown to be dependent on p120-catenin (Vu et al., 2021). While Rac1 is known to function upstream of p190RhoGAP which associates with p120 catenin and influences RhoA activity (Wildenberg et al., 2006), whether Rac1 regulates E-cadherin clustering via p120-catenin is not yet known, but may represent a possible candidate pathway for further investigation. Interestingly, we also note that CA-Rac1 expression appears to give rise to global changes in actin organization, with increased appearance of ventral stress fibers. Altogether these suggest that the actin cytoskeletal architecture of mESCs may depend on low level Rac1 activity and implicate certain RacGAP potentially regulates the unique architecture of the pluripotent states. Conversely, this may also point to a Rac1 downstream effector as being important for stress fibre formation in mESCs. Future molecular dissection will be necessary to delineate the signalling pathways regulating the actin cytoskeleton in these naïve mESCs.

**Figure S1.**
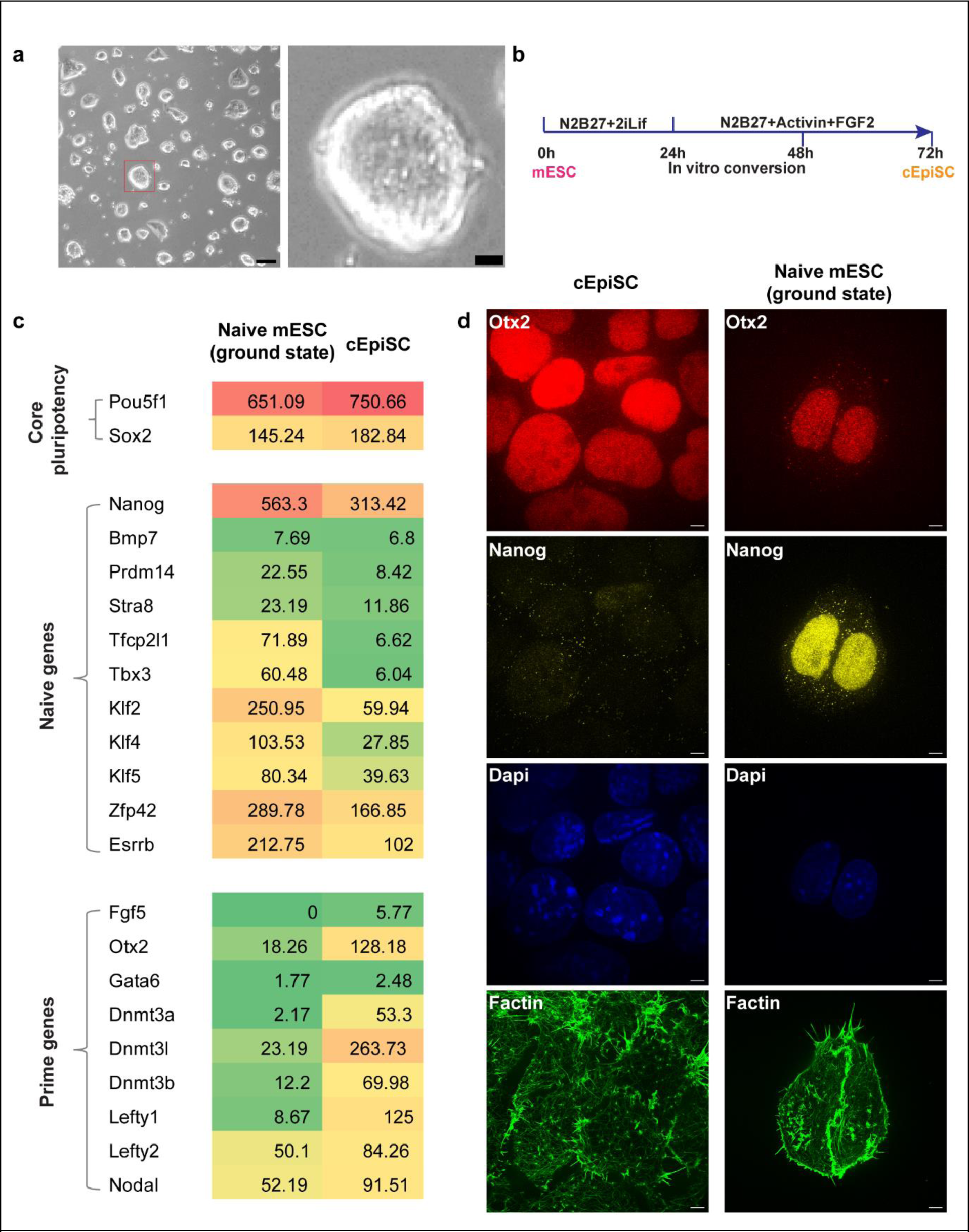
(**a**) Phase contrast images for mESC colonies in 2iLIF ground state at day2. scale bar: 100 μm. Inset image for the red box is at right with scale bar 20 μm. **(b)** *In vitro* protocol for naïve mESC to converted prime EpiSC (cEpiSC) by addition of Activin A and FGF2. **(c)** Heatmap for RNA sequencing RPKM (reads per kilobase of exon per million reads mapped) data for ground state mESCs in 2iLIF and cEpiSCs. **(d)** Corresponding images for Fig. 1c with maximum intensity projection for Otx2, Nanog, and DAPI and F-actin at the ventral plane. Scale bar: 5 µm.

**Figure S2.**
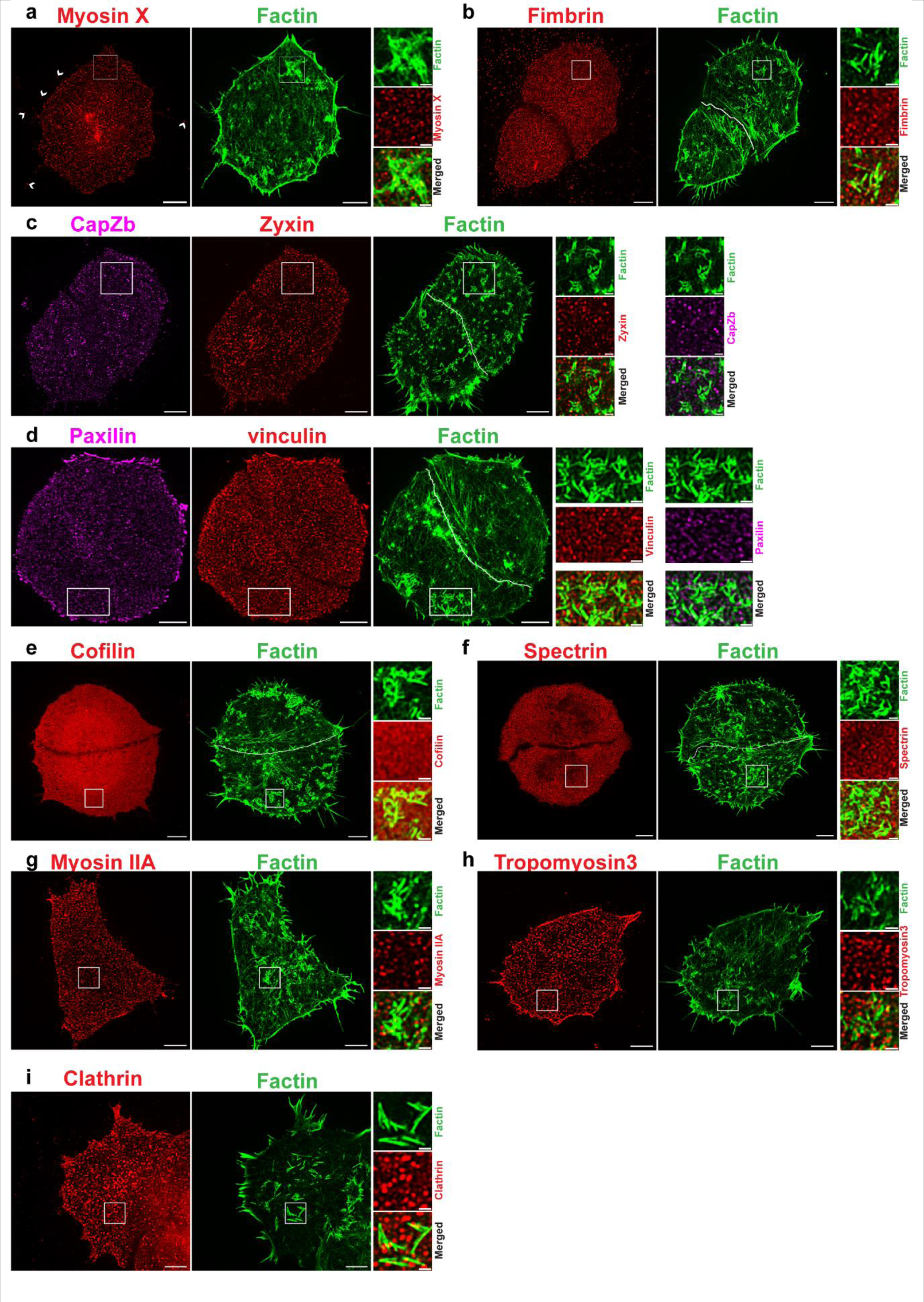
Immunofluorescence images for various actin binding proteins including myosin X (**a**), fimbrin (**b**), capZb and zyxin (**c**), paxilin and vinculin (**d**), cofilin (**e**), spectrin (**f**), myosin IIA (**g**), tropomyosin3 (**h**), and clathrin (**i**). Scale bar: 5 µm. White box highlights the special actin-rich structures of interest, which are not at cell-cell junctions. The white line indicates the intercellular regions in **(b), (c), (d), (e)**, and (**f)** for actin channel. White arrowheads in (**a**) indicate filopodia. Scale bar: 5 µm (inset: 1µm).

**Figure S3.**
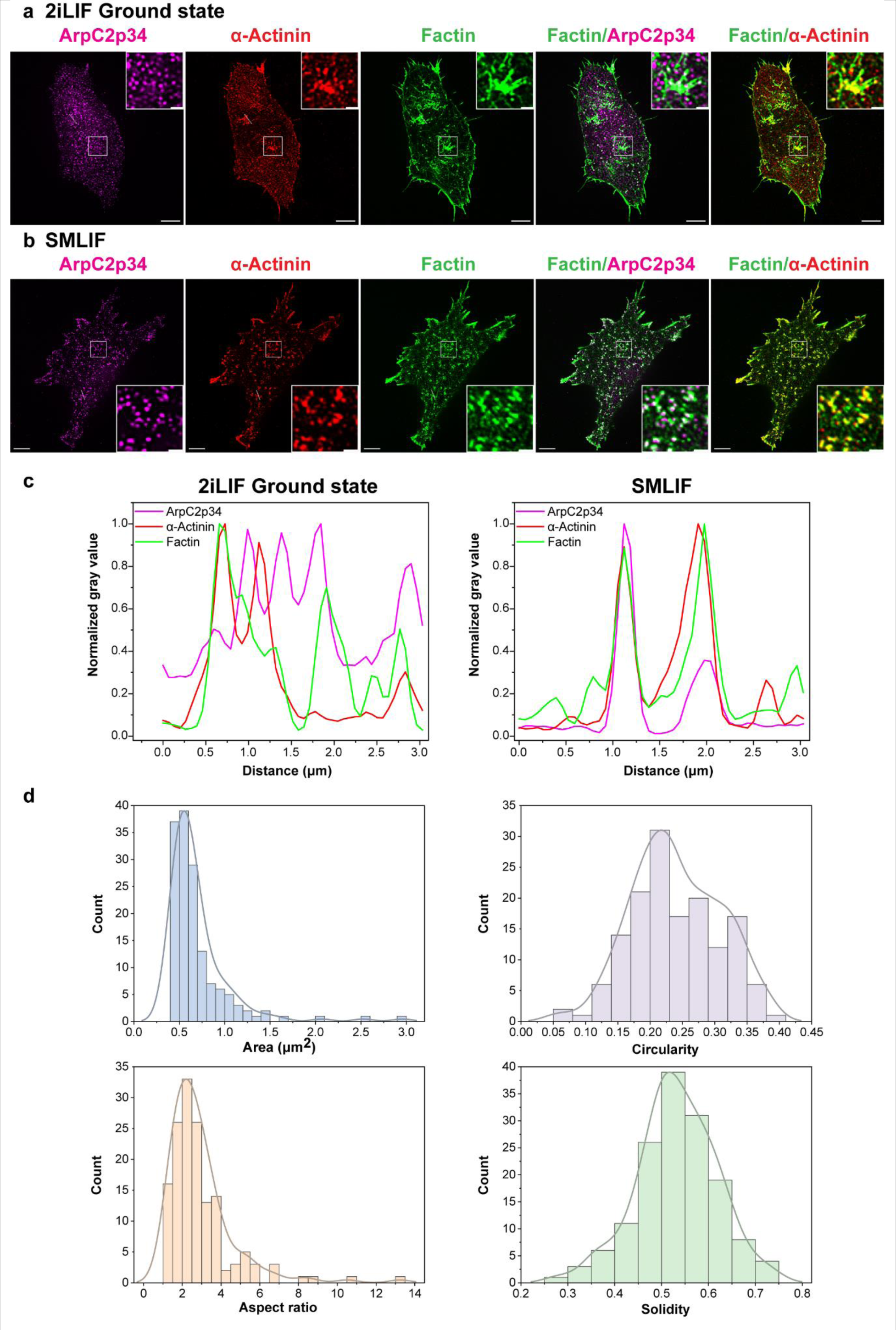
Cadheropodia is different from asters in mESC in SMLIF. (**a**) Confocal micrographs of immunofluorescence staining of ArpC2p34 (magenta), α-Actinin (red), and F-actin (green) for mESC in 2iLIF ground state **(a)** and SMLIF **(b)**. Scale bar: 5 µm (inset: 1µm). **(c)** Line scans of the corresponding channels through the line indicated in image **(a)** and **(b). (d)** Quantification for the cadheropodia in mESC in 2iLIF ground state based on β-catenin channel.

**Figure S4.**
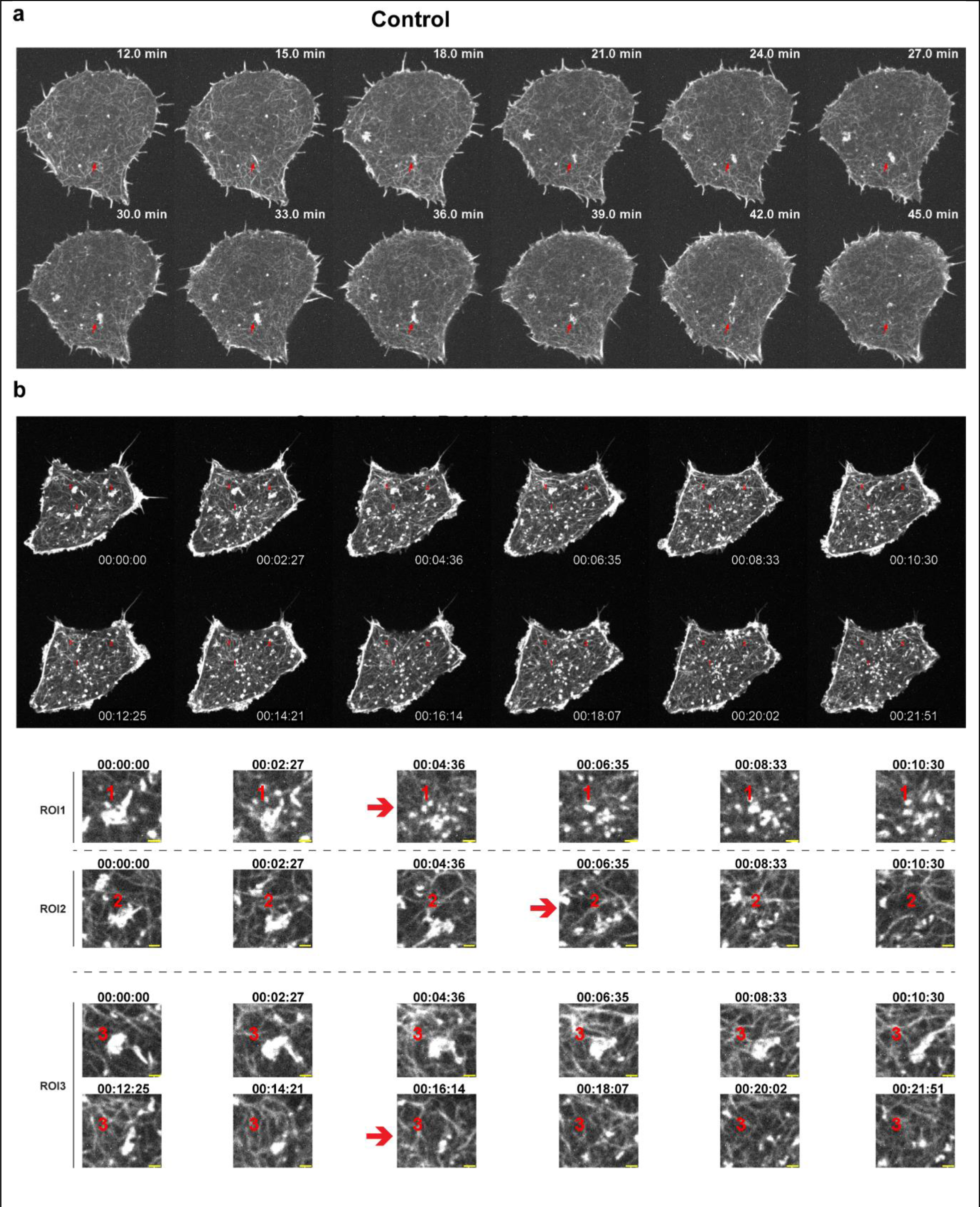
Actin polymerization is required for cadheropodia. Time lapse images for lifeact-Emerald stable cell line of control mESCs in 2iLIF **(a)** and mESCs in 2iLIF after treatment with 0.1 μM cytochalasin D **(b).** Inset scale bar: 1μm.

**Figure S5.**
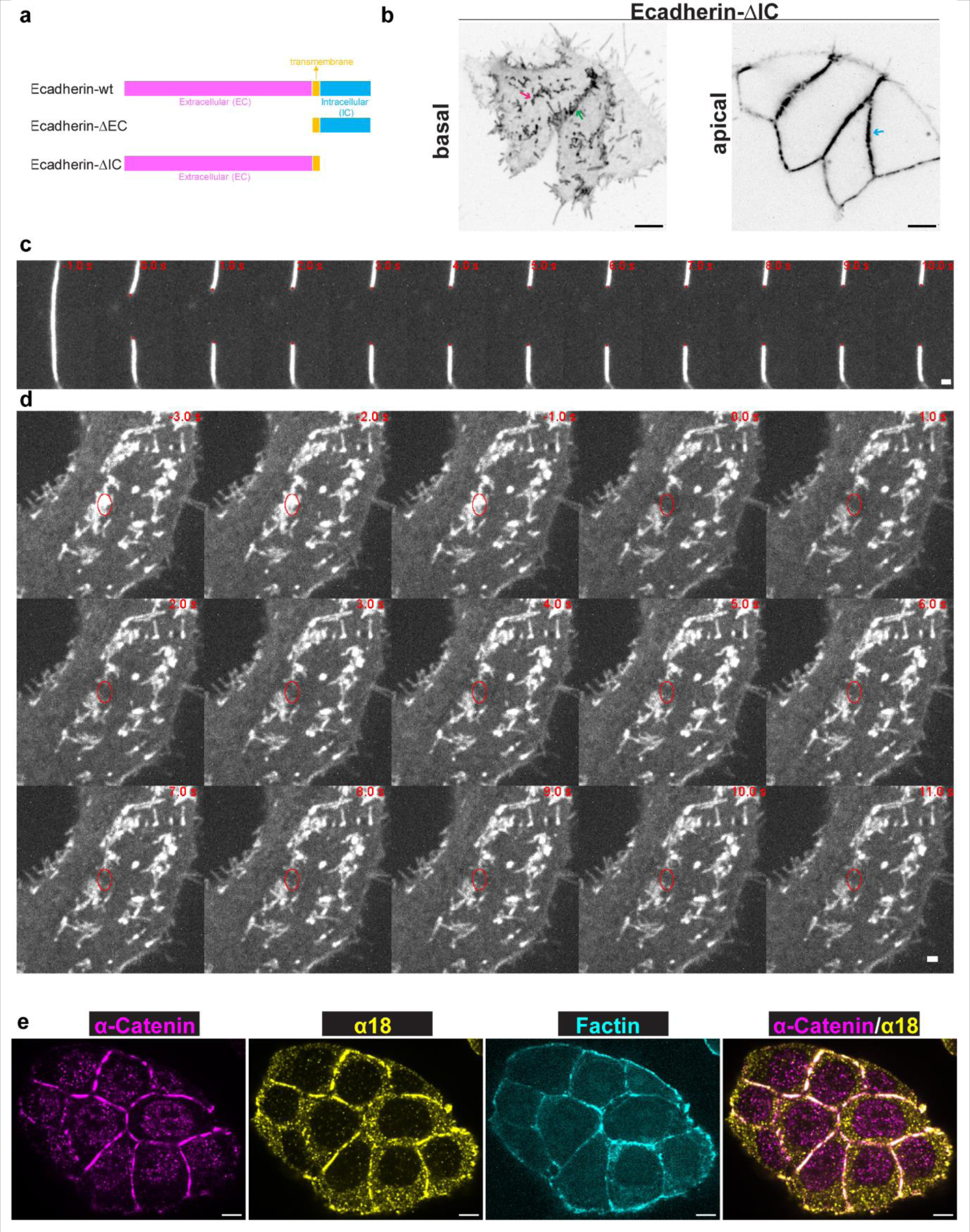
(**a**) Design of wild type and truncated E-cadherin constructs. **(b)** Confocal micrographs for live mESCs in 2iLIF after transfection with E-cadherin without intracellular domain (Ecad-ΔIC) fused with mScarlet for 24h. Scale bar: 5 μm. **(c)** Consecutive images with interval as 1s for E-cadherin-GFP transfected mESC in 2iLIF showing apical junction before and after laser ablation at 0s. Scale bar: 1 μm. **(d)** Consecutive images with interval as 1s for E-cadherin-GFP transfected mESC in 2iLIF showing cadheropodia before and after laser ablation at 0s. Scale bar: 1 μm.**(e)** Confocal micrographs for apical plane of mESCs in 2iLIF after immunofluorescence staining with activated α-catenin (α18), α-catenin, and Factin. Scale bar: 5 μm (inset: 1μm).

**Figure S6.**
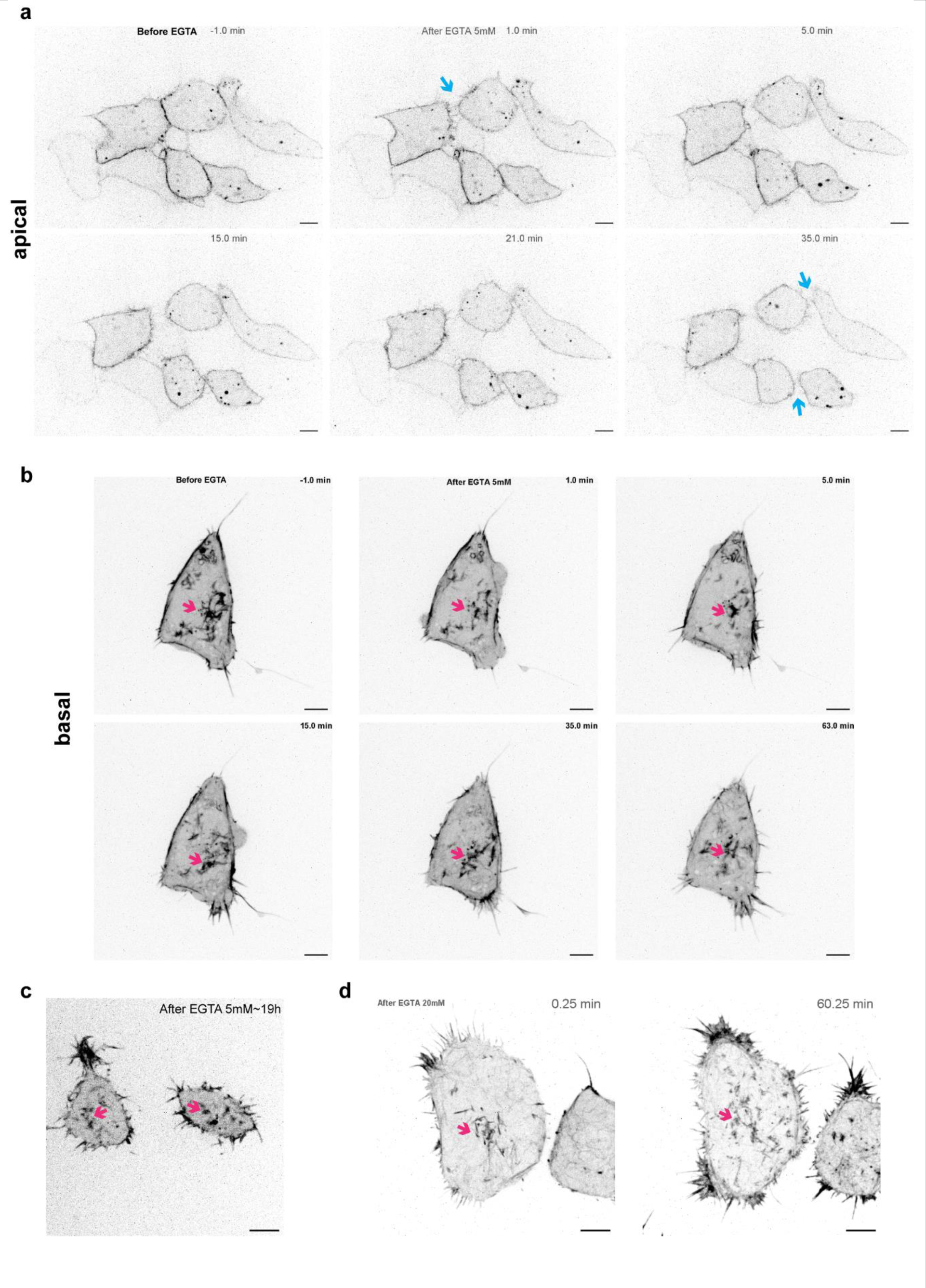
Cadheropodia is more resistant to calcium depletion compared with apical junction. (**a**) Time lapse images exhibit the apical junctions (blue arrows) of mESC in 2iLIF labelled by E-cadherin-mcherry were separated within 35 min after addition of 5 mM EGTA. Scale bar: 5 µm. **(b)** Cadheropodia (magenta arrows) labelled by lifeact-Emerald is still visible in mESC in 2iLIF after addition of 5 mM EGTA for ∼1 h. Scale bar: 5 µm.**(c)** Cadheropodia (magenta arrows) labelled by lifeact-Emerald is still visible in mESC in 2iLIF after addition of 5 mM EGTA for ∼19 h. Scale bar: 5 µm. **(d)** Cadheropodia (magenta arrows) labelled by lifeact-Emerald is still visible in mESC in 2iLIF after addition of 20 mM EGTA for ∼1 h. Scale bar: 5 µm

**Figure S7.**
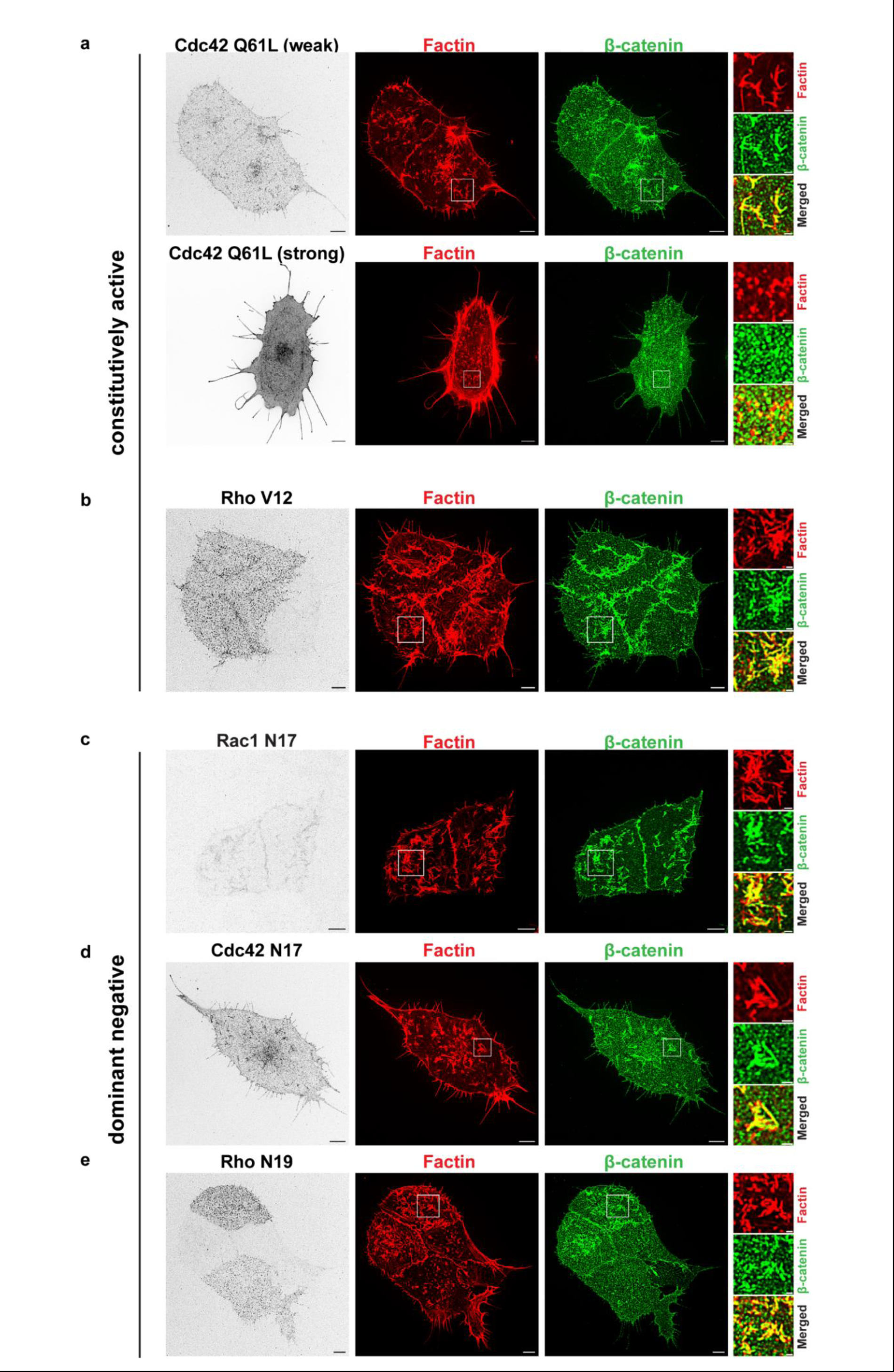
RhoGTPase effect on cadheropodia. Immunofluorescence images for mESC in 2iLIF transfected with constitutively active cdc42 Q61L **(a)**, constitutively active Rho V12 **(b)**, dominant negative Rac1 N17 **(c)**, cdc42 N17 **(d)**, and Rho N19 **(e)**. Scale bar: 5 µm. Note the effect of cdc42 Q61L is expression-dependent, it disassembles cadheropodia and results in long filopodia when the expression is strong.

## Materials and Methods

### Feeder cell culture

Mouse embryonic fibroblast (MEF) were kindly provided by Michael P. Sheetz group and grown in the MEF medium (DMEM with high glucose and GlutaMAX supplement (Gibco™) with 10% embryonic stem-cell qualifiedFBS (Gibco™) and 1 mM sodium pyruvate (Thermo Scientific™)). When MEF cells are confluent, the culture medium was replaced with MEF medium containing 10 µg/ml mitomycin C (Sigma-Aldrich), and incubate for 3 hr at 37°C. After washing twice with D-PBS(-) without Ca and Mg (Nacalai Tesque), cells were trypsinized by 0.25% trypsin/EDTA (Thermo Scientific™) and centrifuged to obtain single cell suspension. Then ∼0.9 million mitomycin C treated MEF cells were seeded on 0.1% gelatin coated 6 cm dish as a feeder layer one day before seeding E14TG2a.

### E14TG2α on feeder cell in SMLIF

ES-E14TG2α (ATCC® CRL-1821™) mouse embryonic stem cells (mESCs) were purchased from ATCC and cultured on mitomycin C-treated feeder initially in the serum containing medium (SMLIF). SMLIF medium consists of DMEM with high glucose and GlutaMAX supplement (Gibco) plus 15% (v/v) Embryonic stem-cell qualified FBS (Gibco), 0.1mM MEM Non-essential amino acids solution (Gibco), 100 U/ml Penicillin and 100 μg/mL Streptomycin (Gibco), 1mM sodium Pyruvate (Gibco), 55 µM β-mercaptoethanol (Gibco). To maintain the pluripotency, 10^3^ units/mL of ESGRO® recombinant mouse LIF is added to the medium and the medium is used within one week. The SMLIF medium is replenished every day and the mESCs were subcultured every 2∼3 days. For subculture, mESCs on feeder cell were trypsinized by 0.05% trypsin/EDTA (Thermo Scientific™), followed by neutralization by serum containing medium without LIF. After centrifugation, cells were suspended in SMLIF and 0.5 million total cells were plated on freshly prepared 6cm feeder cell dish as above.

### E14TG2a on feeder-free in SMLIF

After a few passaging on feeder cell, mESCs were adapted into feeder-free condition but still in SMLIF medium. In the first adaptation, mESCs on feeder cell were trypsinized by 0.05% trypsin/EDTA as above and 0.1 million cells were seeded on 0.1% gelatin coated T25 flask (no feeder cell). Initially, some feeder cells also appear on the flask together with mESC colonies. As MEF attach to the flask stronger than mESCs, treatment with 0.05% trypsin/EDTA for a shorter time such as 1-2min can attach mESCs while leave MEF on the flask. So after subculture for a few times, most feeder cell were removed to obtain a stable feeder-free cultures in SMLIF.

### E14TG2a on feeder-free in 2iLIF

After obtaining E14TG2a feeder-free cultures in SMLIF, they can be further adapted into the ground state 2iLIF condition. The mESCs in feeder-free SMLIF were trypsinized and seeded in SMLIF for the first 24h, followed by media change to 2iLIF. After 2-3 passages in 2iLIF, representative morphology characteristic of ground state mESC with bright domed colonies can be observed. The 2iLIF medium is based on serum-free N2B27. Due to variation in commercial N2 components, N2 was prepared in-house following a previously published protocol (Mulas et al., 2019): 200x N2 consists of 10 mg/mL apotransferrin (Sigma-Aldrich), 1x DMEM/F12 (Sigma-Aldrich), 2.5 mg/mL insulin (Sigma-Aldrich), Bovine Albumin Fraction V (7.5% solution) (Gibco™), 3 μM sodium selenite(Sigma-Aldrich), 1.6 mg/mL putrescine(Sigma-Aldrich), and 1.98 μg/mL progesterone (Sigma-Aldrich). N2B27 is prepared from DMEM/F12 (Sigma-Aldrich), Neurobasal™ Medium (Gibco™), 0.5x B-27™ Supplement (Gibco™), 1 x N2 (as above), 50 μM β-mercaptoethanol (Sigma-Aldrich), 2 mM L-glutamine (Gibco™). N2B27 medium was aliquoted and stored at –80°C. 1 μm PD03 (FGF/ERK inhibitor), 3 μM CHIR99021 (GSK3 inhibitor) and LIF (10^2^ units/mL) were added to N2B27 medium and used within one week. Medium change was performed daily, while cells in 2iLIF condition were subcultured every 2-3 days using accutase on 0.1% gelatin coated 6 well plates with a seeding density of ∼14,000 cells/cm^2^ to ensure the maintenance of colonies with representative morphology.

### *In vitro* conversion of mESCs to mEpiSC

To induce conversion from mESCs to mEpiSC in vitro, mESCs in 2iLIF culture were plated on fibronectin-coated 6-well plates at a density of 2×10^4^ cells/cm^2^ with 2iLIF medium initially. After 24 h, the medium was replaced by N2B27 medium supplemented with 20 ng/mL Activin A and 12 ng/mL FGF2 (RnD Systems). Cells were maintained in the induction media for 2 days. Afterwards, cells were detached by 1 mg/mL collagenase type IV and seeded as small clusters on fibronectin-coated glass-bottom dish (Iwaki) for imaging.

### Transfection

Mouse ESC in 2iLIF was seeded on fibronectin-coated 6 well plates or glass bottom dishes with a diameter of 27mm (Iwaki) at a density of ∼10, 000 cells/cm^2^ overnight and then transfected with 2 μg endotoxin-free plasmids by 4 μl Jetprime reagent following the manufacturer’s protocol (Polyplus). For Ecadherin-wt, Ecadherin-ΔEC, and Ecadherin-ΔIC, after transfection for 24h, live cell samples were imaged directly by spinning disk confocal microscope. For Rho GTPase, cells after transfection for 24h were replated and fixed for immunofluorescence staining.

### Generation of stable cell line

LifeAct-Emerald stable cell line was generated by transfection lifeAct-Emerald into mESCs in 2iLIF on fibronectin coated 6 well plates, followed by Geneticin (G418) 400μg/mL for selection and cell sorting by Sony SH800S Cell Sorter.

### Design of E-cadherin-wt-mscarlet-I and mutants

The E-cadherin-mScarlet-I wild type (wt) and mutants were modified based on the murine-E-cadherin-mCherry, which was a gift from Alpha Yap (Addgene plasmid # 71366; http://n2t.net/addgene:71366; RRID:Addgene_71366) (Han et al., 2014). The mScarlet-I sequence is referred to (Bindels et al., 2017) and carries a point mutation in T74I. The mCherry was replaced with mScarlet-I sequences and fused with murine E-cadherin at C-terminus. The truncated mutants including E-cadherin-mScarlet-I ΔEC (removing extracellular domain 157-709 amino acids) and E-cadherin-mScarlet-I ΔIC (removing intracellular domain 733-884 amino acids) were synthesised by Epoch Life Science, Inc.

### Immunofluorescence staining

Cells were fixed by 4% PFA in 1xPBS for 30 min at room temperature and permeabilized by 0.2% Triton-X in 1xPBS for 15 min. After washing in 1xPBS for three times, the samples were blocked by 4% BSA in 1xPBS for 1 h and incubated with primary antibody in 1% BSA overnight at 4°C. After washing three times in PBS, corresponding secondary antibodies in 1% BSA were added and incubated for 2h at room temperature. To stain cell nucleus, DAPI (330 nM) in 1xPBS was added and incubated for 30 min, followed by washing three times with 1xPBS and imaging freshly by confocal microscopy.

### Spinning Disk Confocal Microscopy and Spinning Disk/instant Structured Illumination Microscopy

Fixed cell samples were imaged on a Nikon Eclipse Ti-E Inverted Microscope (Nikon Instrument, Japan) equipped with a CSU-W1 spinning disk confocal unit (Yokogawa Electric Corporation, Japan), CFI Plan Apochromat ƛ 100x lens, Photometrics Prime 95B Scientific CMOS camera, and 405 nm, 488 nm, 561 nm, 642 nm lasers. For Spinning Disk/instant Structured Illumination Microscopy (SD-SIM) imaging, the instant Structured Illumination Microscopy module (Live-SR, Gataca Systems, France) in the microscope system was used, which provides a pixel size of 65.87 nm (xy). For live cell imaging, temperature controller (LCI CU-501) and CO2 mixer (LCI FC-SN) were used to maintain the imaging chamber at 37 °C and 5% CO_2_. The 2iLIF medium is used for live cell imaging.

For pharmacological perturbation by Cytochalasin D or Blebbistatin, Lifeact-Emerald stable cell line was seeded on fibronectin-coated glass bottome dish (Iwaki), and live cell imaging was started after 3-6h seeding. Timlapse was taken before and after treatment with 0.1 μM Cytochalasin D (Invitrogen™) or 10 μM *para*-nitro Blebbistatin (Cayman Chemical).

For treatment with EGTA, Lifeact-Emerald stable cell line was transfected with E-cadherin-mcherry for 24h. After replating for 5h-24h, timlapse was conducted before and after treatment with 5, 10, or 20 mM EGTA (Sigma-Aldrich).

For treatment with Decma antibody, mESCs in 2iLIF were transfected with E-cadherin GFP for 24h. After replating for 4-24h, timlapse was conducted before and after treatment with 10 μg/mL Decma antibody (Sigma-Aldrich).

### Laser ablation

The apical junction and cadheropodia were selectively ablated by UV laser nanoscissor performed on a Nikon Eclipse Ti inverted microscope (Nikon Instrument, Japan) equipped with a CSU-W1 spinning disk confocal unit (Yokogawa Electric Corporation, Japan), a LED-based epifluorescence excitation source (X-lite 110 LED), a laser combiner (405 nm, 488 nm, 561 nm, and 642 nm solid-state lasers, and 355nm pulsed ablation laser (GATACA systems), Photometrics Prime 95B sCMOS camera (Photometrics, USA), 100x ablation lens (Nikon), stage top incubator and CO2 mixer (LCI). The iLas Pulse system (GATACA Systems) in the Modular GATACA software was used to control the ablation area. After calibration, time-lapse confocal images were acquired every 1 s, starting from 2 frames prior to the ablation until 30 frames post-ablation.

### ZEISS Lattice Lightsheet 7 imaging

Mouse ESCs in 2iLIF were transfected with E-cadherin-GFP for 24h and replated on AlphaPlus 35mm dish for overnight. Then the sample was imaged by ZEISS Lattice Lightsheet 7 equipped with PCO Edge camera, 13.3x / Numerical Aperture 0.4 illumination objective lens, 44.83× / Numerical Aperture 1.0 detection objective, and generating a lightsheet with Sinc3 30×1000 (30um of beam length, 1000nm of beam thickness). The fluorescence was excited by 488 nm and emission is collected at 500-588 nm. The raw data was deskewed with cover glass transformation, which was then further processed with deconvolution using a restorative algorithm. To be more precise, constrained iterative algorithm was used where 12 iterations were performed under the following parameters: Strength: 8, Regularization: Zero Order, Likelihood: Poisson and PSF: Theoretical PSF.

### RNA-SEQ

Total RNA were extracted from mESCs in 2iLIF and cEpiSC after conversion by RNeasy® Plus Mini Kit (Qiagen) following the manufacturer’s protocol. Then the RNA samples were sent to BGI Genomics (Shenzhen, China) for whole genome RNA sequencing using BGISEQ platform, averagely generating about 4.49G Gb bases per sample. The average mapping ratio with reference genome is 96.46%, the average mapping ratio with gene is 88.98%; 18032 genes were identified.

### Image analysis

To quantify cadheropodia in mESC in 2iLIF fixed at 6h, immunofluorescence staining labelling F-actin and β-catenin were taken on CSU-W1 spinning disk confocal microscope. The β-catenin channel images were then background-subtracted using ImageJ, with local thresholding by Bernsen method. ROI defining the cell cortex were then manually defined, excluding the cell edge. Subsequently, cadheropodia structures within each ROI were quantified using “Analyze Particle” function, whereby structures smaller than 0.4 μm^2^ and circularity >0.4 were removed.

To quantify the enrichment of Ecad-wt and Ecad-ΔEC (Fig 4), the numbers of cells with Ecad-wt or Ecad-ΔEC mScarlet-I localization at cadheropodia, basal CCJ, and apical CCJ were counted manually and divided by the total number of analyzed cells to obtain the corresponding percentages. The percentages of Ecad-ΔEC were then divided by the corresponding values of Ecad-wt for normalization for every independent batch.

To analyse the percentage of cells with cadheropodia in control and Rac1 V12 transfected cells, the numbers of cells with cadheropodia in different groups were counted manually based on shape and colocalization of β-catenin and F-actin, which were divided by the total analysed cells with appropriate transfection signals to obtain the corresponding percentages.

### Statistics

Quantitative data were analysed for normality using a Shapiro-Wilk normality test. Data with a Gaussian distribution were analysed using a two-tailed unpaired Student’s t-test (two groups). Significant differences in the variance were taken into account using a Welch’s correction. Data that did not have a Gaussian distribution were analysed using a Mann-Whitney U-test (two groups).

## Author Contributions

S.L. and P.K. designed the project and wrote the manuscript. Y.M. helped design E-cadherin-mScarlet-i wt, ΔEC, ΔIC, and contribute reagents.

## Acknowledgements

We gratefully acknowledge Diego Pitta de Araujo for generating the graphical illustrations, and thank Ong Hui Ting for helping some of the image analysis.

## Antibodies

| Antibodies | Source | Catalogue # | Dilution |
| --- | --- | --- | --- |
| Mouse anti-oct4 | Sigma-Aldrich | MAB4419 | 1/200 |
| Rabbit anti-nanog | Abcam | ab80892 | 1/200 |
| Goat anti-KLF4 | R&D Systems | AF3158 | 1/200 |
| Goat anti-Otx2 | R&D Systems | AF1979 | 1/200 |
| Rabbit anti-a-catenin | Abcam | ab51032 | 1/200 |
| Rabbit anti-β-catenin | Abcam | ab2365 | 1/200 |
| Mouse anti-β-catenin | BD Biosciences | 610154 | 1/200 |
| Mouse anti-E-cadherin | BD Biosciences | 610181 | 1/200 |
| Rabbit anti-p120 | Santa Cruz | sc-13957 | 1/100 |
| Rabbit anti-paxilin | Santa Cruz | sc-5574 | 1/200 |
| Mouse anti-a-actinin | Sigma-Aldrich | A5044 | 1/200 |
| Rabbit anti-ARPC2p34 | Sigma-Aldrich | 07-227 | 1/200 |
| Mouse anti-vinculin | Sigma-Aldrich | V9131 | 1/200 |
| Mouse anti-zyxin | Abcam | ab58210 | 1/200 |
| Rabbit anti-capzb | Thermo Fisher | 25043-1-AP | 1/200 |
| Rabbit anti-cofilin, | Abcam | ab11062 | 1/200 |
| Mouse Anti-β-Spectrin II | BD Biosciences | 612563 | 1/200 |
| Rabbit anti-clathrin | Abcam | ab21679 | 1/200 |
| Rabbit anti-myosin X | Sdix | 22430002 | 1/200 |
| Rabbit anti-fimbrin | Abcam | ab236976 | 1/200 |
| Rabbit anti-Tropomyosin 3 | Abcam | ab180813 | 1/200 |
| Rabbit anti- myosinIIA | Cell Signaling Technology | 3403S | 1/50 |
| Rat anti-α-catenin (active),<br>α18 | Akira Nagafuchi | | 1/500 |

## Plasmids

| Plasmids | Sources |
| --- | --- |
| mCherry-Lifeact-7 | Addgene #54491 |
| mEmerald-Lifeact-7 | Addgene #54148 |
| E-cadherin-GFP | Addgene #67937 |
| E-cadherin-mcherry | Addgene #71366 |
| E-cadherin-mScarlet-i wild type (wt) | Custom-synthesized by Epoch Life Sciences, Inc. , USA |
| E-cadherin-mScarlet-I-ΔEC | Custom-synthesized by Epoch Life Sciences, Inc. , USA |
| E-cadherin-mScarlet-I-ΔIC | Custom-synthesized by Epoch Life Sciences, Inc. , USA |
| Rac V12-BFP | Michael P. Sheetz (MBI, Singapore) |
| Rac N17-BFP | Michael P. Sheetz (MBI, Singapore) |
| pcDNA Cdc42 Q61L-EGFP | Clare M. Waterman (NIH, USA) |
| Cdc42 N17-EGFP | Michael P. Sheetz (MBI, Singapore) |
| RhoA N19-EGFP | Michael Sheetz (MBI, Singapore) |
| RhoA V12-EGFP | Michael Sheetz (MBI, Singapore) |

